# Gastrointestinal parasite community in the critically endangered West African lion

**DOI:** 10.1101/2020.06.09.136176

**Authors:** Sofia Kruszka, Nyeema C. Harris

## Abstract

Large carnivores of Africa, such as lions (*Panthera leo*), suffer from prey depletion and habitat fragmentation, that possibly impact the composition of the gastrointestinal parasite community. West African lions are particularly important, as this population is critically endangered and yet little is known of their gastrointestinal parasite community, which can reflect the health and resilience of the host population. From fecal samples collected in the W-Arly-Pendjari (WAP) transboundary protected area complex in Burkina Faso and Niger, we identified 309 oocysts of at least five different species using fecal flotation and sedimentation tests. We also compared these gastrointestinal parasites to other results from surveys of lions from Southern and East African regions and found similar taxa to previous surveys, but lower species richness across West African samples.

---

Large carnivores are in jeopardy (Dirzo et al., 2014; Ripple et al., 2014). Reductions in their population sizes and geographic ranges have broad ecological ramification in structuring communities and ecosystem processes through consumptive and non-consumptive pathways (Schmitz et al., 2010; Abdala-Roberts et al., 2019). One such consequence includes variation in parasite communities emergent from changing host densities that affect disease dynamics and health attributes (Colwell et al., 2012; Hatcher et al., 2012; DeCandia et al., 2019). For example, fragmentation and other land degradations coupled with pollution can alter micro- and macroparasite diversity (Altizer et al., 2003; Chakraborty et al., 2019), which in turn impairs the host’s ability to digest nutrients and defend itself against more harmful pathogens (Amato et al., 2013). Therefore, we underscore that studying parasite composition and diversity are important for assessing health particularly in threatened populations.

Studies of parasites in threatened carnivores provide essential ecological information that can aid in promoting their recovery and persistence. In the endangered Iberian lynx (*Lynx pardinus*), *Ancylostoma tubaeforme* was significantly more prevalent in juveniles than adults in Doñana National Park in Southwest Spain, and the authors highlight investigating the clinical consequences of hookworm burdens (Vicente et al., 2004). Intestinal parasites detected in feces or other parasites found on the host can reflect nutritional requirements and dietary preferences. A gastrointestinal survey of scat from endangered Croatian wolves (*Canis lupus*) indicated consumption of fish, wild boar (*Sus scrofa*), roe deer (*Capreolus capreolus*), and deer (*Cervus elaphus*) based on the detection of *Diphyllobothrium* sp. and *Opisthorchis* sp. helminths and a high prevalence of *Sarcocystis* sp. protozoa (Hermosilla et al. 2017). In a survey of ectoparasites for endangered black-footed ferrets (*Mustela nigripes*) in South Dakota, Harris and colleagues (2014) observed parasites associated with common prey species including prairie dogs (*Cynomys* sp.).

Africa’s iconic apex predator, the African lion (*Panthera leo*), faces a tenuous future with persistent threats from prey depletion, human conflict, and habitat loss (Bauer et al., 2015). As such, different populations may harbor a unique parasite community with varying implications for the host’s health and spatial patterns of biodiversity, more broadly. To date, several studies have investigated intestinal parasites found in feces from free-ranging lions in East and South Africa. (Müller-Graf 1995; Berensten et al. 2012; Lajas et al., 2015). Here, we present the first survey of the gastrointestinal parasite community for lions in West Africa. This critically endangered population now occupies only 1% of historic range and continues to decline (Henschel et al., 2014). Our work: 1) validates methods to confirm host identity from fecal samples through kit and primer testing; 2) describes the gastrointestinal parasite community of lions in West Africa; and 3) conducts regional comparisons of the gastrointestinal community across lion populations through literature searches. Overall, we expect a parasite community comprised of generalist parasites for West African lions due to their lower population sizes and increased vulnerability to extinction.

Specifically, we investigated the gastrointestinal parasites of lions in the largest West African population located in the W-Arly-Penjari (WAP) transboundary protected area complex. WAP is a UNESCO World Heritage site and the largest protected area complex in West Africa, comprising 26,515 km^2^ across Niger, Benin, and Burkina Faso. Lions coexist with other large carnivores, namely spotted hyenas (*Crocuta crocuta*) and leopards (*Panthera pardus*) and a myriad of ungulate prey in a network of national parks and hunting concessions (Mills et al., 2020). However, human pressures persist, as livestock grazing is pervasive, which may exacerbate disease risks for sympatric wildlife (Harris et al., 2019).

From 2016-2018, we collected feces presumed to be lion in WAP and preserved samples in ethanol (EtOH), N,N-diethyltryptamine (DET) buffer, or RNAlater for subsequent molecular confirmation and analysis. We extracted DNA by testing protocols from three different manufacturer kits: Qiagen© QIAmp (MoBio) PowerFecal DNA Kit, MP© FastDNA SPIN Kit for Feces, and Qiagen© QIAmp DNA Stool Mini Kit. DNA were amplified through PCR using three different primers for target genes: 16S (Tessler et al., 2017), 12S (Kitano et al., 2007), and CytB (Kocher et al., 1989; Farrell et al., 2000). Samples were then matched to known sequences in the NCBI database with 98% or higher identity and 90% or higher query cover being confirmed as African lion. We used 2g of feces for fecal flotation tests using a ZnSO4 solution diluted to a specific gravity of 1.2 with centrifugation to isolate and observe lighter intestinal parasites such as nematodes and cestodes. Additionally, to detect heavier parasites such as trematodes, we used sedimentation test in ZnSO4 solution without centrifugation. We used measurements obtained through microscopy and morphology described by Zajac et al (2012) to complete identifications.

We conducted a literature search through Web of Science to compile known gastrointestinal parasites reported from wild, free-ranging African lions. Keywords included: “gastrointestinal”, “helminth”, “macroparasite”, “gut”, “parasite”, “disease”, “*Panthera leo*”, and “African lion”. We filtered search results to remove any experimental manipulations or studies occurring in captive populations. From resultant papers, we also conducted backwards and forward searches to identify any additional gastrointestinal papers found for African lions.

Of the three extraction kits tested, we recommend the Qiagen © QIAmp DNA Stool kit, which required the least amount of sample and was relatively easy to train personnel to implement. Of the three universal genes, the 16S primers produced the highest quality amplicons, allowing for the highest percent of host identifications for the Qiagen © QIAmp DNA Stool kit (85%). We found CytB performed the worst in amplification of lion samples across all kits with only 11-22% confirmation of lion samples.

We identified 309 oocysts of at least five different gastrointestinal parasite species in 11 confirmed samples from critically endangered West African lions in WAP (Figure 1). All parasites identified were previously reported in South and East African lion populations, resulting in no new host records (Table 1). We found one egg that morphologically resembled two attached *Sarcocystis* sp. sporocysts based on the egg dimensions of 17.9 × 14.5 micrometers (Zajac et al., 2012). However, because it is not common to find the sporocysts still attached after leaving the host (Wassermann et al., 2017), we state as unknown (Figure 1f), though *Isospora* sp. could also be a plausible identification.

**Figure 1.**
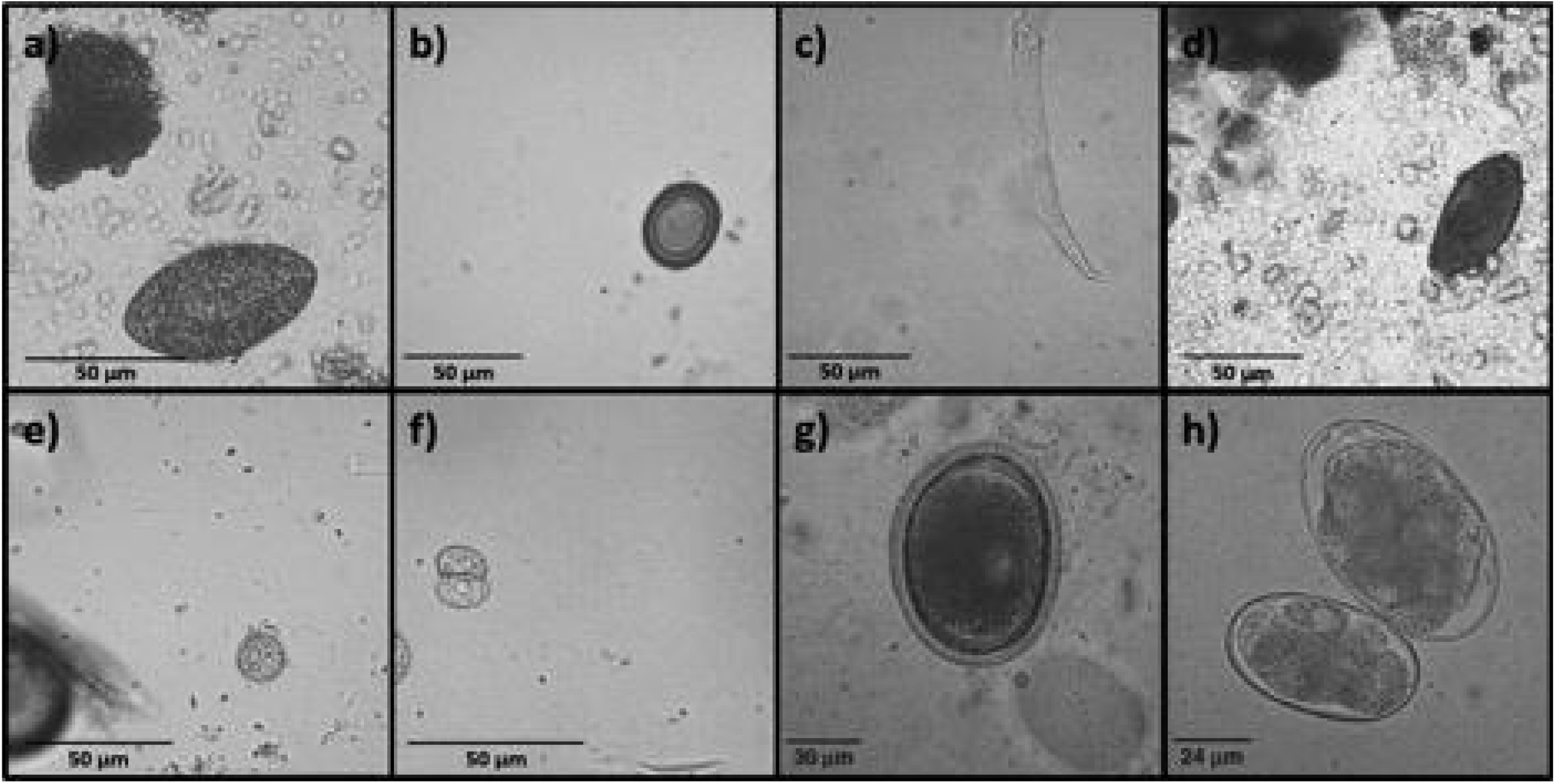
Species found in West African lion samples: a) *Spirometra sp*., b) *Taenia sp*., c) *Aelurostrongylus* larva, d) *Trichuris sp*., e) *Isospora sp*. oocyst, f) Unknown oocyst, possibly *Sarcocystis sp*. Species commonly reported in our lion populations: g) *Toxocara sp*., h) *Ancylostoma sp*. (g-h photo credit: Zajac et al., 2012).

**Table 1.** Gastrointestinal parasite species identified in East, Southern, and West African lion samples. The percentage values represent the species relative abundance from the samples in this study. These species relative abundance calculations exclude two samples: one with too many *Spirometra sp*. to count, and one with too many *Isospora sp*. to count.

| Species | East Africa | Southern Africa | West Africa |
| --- | --- | --- | --- |
| **Protozoa** |  |  |  |
| *Eimeria* sp. | X |  |  |
| *Isospora felis* | X | X |  |
| *Isospora rivolta* | X |  |  |
| *Isospora* sp. | X* |  | 1% |
| *Sarcocystis* sp. | X |  |  |
| *Toxoplasma-like* sp. | X |  |  |
| *Toxoplasma gondii* |  | X |  |
| *Giardia* sp. | X |  |  |
| **Nematoda** |  |  |  |
| *Aelurostrongylus* sp. | X* | X* | 19% |
| *Aelurostrongylus abstrusus* |  | X |  |
| *Ancylostoma* sp. | X |  |  |
| *Ancylostoma braziliense* |  | X |  |
| *Ancylostoma paraduodenale* | X |  |  |
| *Anclystoma tubaeforme* | X* |  |  |
| *Capillaria* sp. | X |  |  |
| *Dirofilaria sudanensis* |  | X |  |
| *Gnathostoma spinigerum* |  | X |  |
| *Habronema* sp. | X |  |  |
| *Lagochilascaris major* | X |  |  |
| *Ollulanus tricuspis* | X |  |  |
| *Physaloptera* sp. | X |  |  |
| *Physaloptera praeputialis* | X |  |  |
| Spirurida | X |  |  |
| *Toxascaris leonina* | X |  |  |
| *Toxocara* sp. | X* | X* |  |
| *Toxocara cati* | X |  |  |
| *Toxocara canis* | X |  |  |
| *Trichinella spiralis* |  | X |  |
| Trichostrongylidae | X |  |  |
| *Trichuris* sp. | X |  | 1% |
| *Uncinaria stenocephala* |  | X |  |
| **Acanthocephala** |  |  |  |
| Acanthocephala | **X** |  |  |
| **Arthropoda** |  |  |  |
| *Linguatula sp.* |  | **X** |  |
| *Linguatula serrata* |  | **X** |  |
| *Linguatula nuttalli* |  | **X** |  |
| **Platyhelminthes** |  |  |  |
| *Spirometra sp.* | **X\*** | **X\*** | **57%** |
| *Spirometra okumurai* | **X** |  |  |
| *Anoplocephalidae* | **X** |  |  |
| *Taenia sp.* | **X\*** | **X\*** | **14%** |
| *Taenia regis* (T. bubesei) | **X** | **X** |  |
| *Taenia gonyamai* | **X** |  |  |
| *Echinococcus sp* | **X\*** |  |  |
| *Echinococcus granulosus* | **X** | **X** |  |
| *Echinococcus felidis* | **X** | **X** |  |
| Taeniidae | **X\*** | **X\*** |  |
| *Schistosoma mattheei* |  | **X** |  |
| *Paramphistomum sp.* |  | **X** |  |
| *Spirometra ranarum* | **X** |  |  |

We found twelve studies surveying the gastrointestinal parasite diversity of wild, free-range lions from six sites in East Africa – Serengeti National Park, Selous Game Reserve in Tanzania, Tarangire National Park in Tanzania, Nairobi National Park in Kenya, Queen Elizabeth National Park in Uganda, and South Luangwa National Park in Zambia – and two different sites in Southern Africa – Niassa National Reserve in Mozambique, and Kruger National Park in South Africa (Ortlepp, 1937; Pitchford et al., 1974; Rodgers, 1974; Young, 1975; Bwangamoi et al., 1990; Müller-Graf, 1995; Müller-Graf et al., 1999; Bjork et al., 2000; Huettner et al., 2009; Berensten, 2012; Kavana et al., 2015; Lajas et al., 2015; Eom et al., 2018). Most studies surveyed fecal samples; however, one study used post-mortem analysis. The average number of individual lion samples reported per study was 29 with a number of studies sampling from only one individual and others that sampled from more than forty. Several parasites that were commonly found in South and East African lions were not identified in our West African lion samples, including *Toxocara* sp. and *Ancylostoma* sp. (Müller-Graf, 1995; Bjork et al., 2000; Berentsen et al., 2012; Lajas et al., 2015).

Our data support the expectation that smaller populations including species of conservation concern will have a less diverse parasite assemblage (Altizer et al., 2007, Harris et al., 2010). However, we recognize more samples are needed from West Africa to make more definite comparisons. As a host population decreases in size, the richness of specialist parasites may also decline, leading to a greater proportion of generalist parasites in a host population (Dunn et al., 2009, Harris et al., 2014). This pattern is unfortunate for threatened species, as generalist parasites tend to be more deleterious to small populations (Stringer et al., 2014). Additionally, there are possibly negative implications for the wild carnivore hosts that follow macroparasite infection (Seltmann et al., 2019). Two studies reported pneumonia or bronchitis in European wildcats (*Felis silvestris silvestris*) infected with species of the lungworm genus *Aelurostrongylus* (Veronesi et al., 2016; Stevanović et al., 2019). Given the present threat of extinction for West African lions, future works should determine which parasites found in lion populations have fitness consequences as well as investigate the gut microbiome, given implications on immune and metabolic function (Ferreira et al., 2019; Karmacharya et al., 2019).

In conclusion, we provide specific methodological suggestions for host confirmation from scat for a critically endangered apex carnivore. Our work reveals relatively lower diversity of parasites in West African lions compared to populations in other regions throughout the continent. Comparative studies of gastrointestinal communities and their impact on host fitness will aid scientists and managers alike to assess the overall health of lion populations in Africa, and identify the regions in which conservation efforts and population management are most needed.

## Acknowledgements

We thank the Applied Wildlife Ecology Lab for assistance with protocol development for labwork and parasite identification especially B. Aguilar and M. Clayson. We extend our sincerest appreciation to all park managers and the field team including I. Gnoumou and Y.I. Abdel-Nasser that assisted with sample collection in the field. We also thank administrators in Ministries of Environment in Burkina Faso (OFINAP & DGEF) and Niger (DGE/EF) especially B. Doamba and Y. Harissou for permitting and assisting with field management as well as private concessionaires in Burkina Faso for wildlife management efforts and access to properties in WAP. We also acknowledge University of Michigan (UM) African Studies Center – STEM initiative, UM Office of Research, the German Society of Mammalian Biology, and the Detroit Zoological Society for financial support.

## Author contributions

SK completed all lab analysis and wrote the paper. NCH conceptualized and managed project, secured funded, led field efforts and data collection, and edited manuscript.

